# Cellular pathways during spawning induction in the starlet sea anemone *Nematostella vectensis*

**DOI:** 10.1101/2021.03.16.435650

**Authors:** Shelly Reuven, Mieka Rinsky, Vera Brekhman, Assaf Malik, Oren Levy, Tamar Lotan

## Abstract

In cnidarians, long-term ecological success relies on sexual reproduction. The sea anemone *Nematostella vectensis*, which has emerged as an important model organism for developmental studies, can be induced for spawning by temperature elevation and light exposure. To uncover molecular mechanisms and pathways underlying spawning, we characterized the transcriptome of *Nematostella* females before and during spawning induction. We identified an array of processes involving numerous receptors, circadian clock components, cytoskeleton, and extracellular transcripts that are upregulated upon spawning induction. Concurrently, processes related to the cell cycle, fatty acid metabolism, and other housekeeping functions are downregulated. Real-time qPCR revealed that light exposure has a minor effect on expression levels of most examined transcripts, implying that temperature change is a stronger inducer for spawning in *Nematostella*. Our findings reveal the mechanisms that may enable the mesenteries to serve as a gonad-like tissue for the developing oocytes and expand our understanding of sexual reproduction in cnidarians.

**Summary statement:** Analysis of transcriptional changes during spawning induction in *Nematostella vectensis*, revealed upregulation of processes related to signal perception and cytoskeleton rearrangement and downregulation of fatty acid metabolism and housekeeping processes.

## Introduction

Sexual reproduction is the predominant mode of procreation in almost all eukaryotes, from fungi and plants to fish and mammals. It generates the conditions for sexual selection, which is a powerful evolutionary force driving morphological, physiological, and behavioral changes in many species (Bai, 2015; Heitman, 2015). Sex is thought to have arisen once and to have been present in the last eukaryotic common ancestor (Goodenough and Heitman, 2014; O’Malley et al., 2019; Speijer et al., 2015); therefore, it is an important trait in evolutionary biology. An informative group of animals for studying evolution are cnidarians, as they are basal metazoans and considered to be a sister group of the Bilateria (Martindale et al., 2004). Within cnidarians, the anthozoans, a group that includes corals and sea anemones, are among the morphologically simplest extant eumetazoans. Although they can reproduce asexually, their long-term ecological success relies on sexual reproduction (Ball et al., 2004; Miller and Ayre, 2004).

The most well-known sexual reproduction phenomenon in cnidarians is the mass spawning of corals. In this process, which has been observed worldwide, individuals from the same species synchronously release their eggs and sperm to the water column, where fertilization occurs (Baird et al., 2009; Hagman et al., 1998; Harrison et al., 1984). The synchronized spawning is postulated to be controlled by the circadian clock mechanism, which is based on transcription–translation feedback loops and maintains a free-running period of approximately 24 hours (Dunlap, 1999). This allows organisms to anticipate and adjust to daily and seasonal environmental cycles (Pittendrigh, 1993; Shearman et al., 2000). The circadian clock obtains environmental cues such as sea temperature, lunar phase and the daily light cycle (Babcock et al., 1986; Babcock et al., 1994; Harrison et al., 1984) that activate internal cascades, eventually leading to spawning (Baird et al., 2009; Kaniewska et al., 2015; Levy et al., 2007; Rosenberg et al., 2017). Orthologs of circadian genes, such as *Bmal*, C*lock, Cry1* and *Cry2* and *Timeless*, have been identified in most cnidarian genomes (Reitzel et al., 2013; Shoguchi et al., 2013; Vize, 2009).

The starlet sea anemone, *Nematostella vectensis*, has become an important model organism for developmental and evolutionary studies, particularly for comparative studies of the conservation of developmental processes and signaling pathways in cnidarians versus bilaterians (Layden et al., 2016; Rentzsch and Technau, 2016). Its genome has been sequenced (Putnam et al., 2007) and spawning can be induced in the lab throughout the year by temperature elevation and light exposure (Fritzenwanker and Technau, 2002; Hand and Uhlinger, 1992; Stefanik et al., 2013). *Nematostella* polyps can be either male or female and can reproduce asexually by transverse fission or sexually by releasing gametes into the water column, where fertilization occurs (Darling et al., 2005; Genikhovich and Technau, 2009). As most anthozoans, *Nematostella* lack real ovaries and the mesenteries serve as gonads, where oocytes develop and mature (Layden et al., 2016), leading to changes in tissue shape (Levitan et al., 2015). The oocytes first appear at the base of the mesenterial gastrodermis and develop asynchronously into the mesoglea while keeping contact with the gastrodermis through the trophonema (Eckelbarger et al., 2008; Moiseeva et al., 2017). However, the natural sexual reproduction of *Nematostella* is still unexplored and it is currently unknown whether it is controlled by the circadian clock or by other mechanisms (Hand and Uhlinger, 1992).

Here, we applied comparative transcriptomic methods to reveal the potential cellular and molecular pathways underlying spawning in *Nematostella*. By characterizing the transcriptome of *Nematostella* females before and during the different stages of spawning induction, we identified an array of upregulated processes involving numerous receptors as well as extracellular matrix (ECM) organization and regulation of actin, along with downregulated processes related to cell cycle, fatty acid metabolism and other housekeeping functions. The regulation of these processes sheds light on the mechanisms that may enable the mesenteries to serve as a gonad-like tissue for the developing oocyte.

## Materials and Methods

### *Nematostella* cultivation

*Nematostella* were kept in artificial seawater (12.5 ppt, pH 8.0, Red Sea, Israel) at 18°C in the dark. They were fed five times a week with freshly hatched brine shrimps (*Artemia salina*) with weekly seawater changes. Spawning was controlled and induced as described (Fritzenwanker and Technau, 2002). In short, on the day before spawning induction, the anemones were not fed, and spawning was conducted in an incubator under white light by raising the temperature from 18°C to 25°C for 12 h.

### RNA-seq experimental design

For the RNA-seq experiment, 25 well-fed female anemones were collected. As a control, 5 individual anemones were placed in a dark incubator at 18°C, sampled before spawning induction, and immediately snap-frozen in liquid nitrogen. Spawning was then induced as described above. At 1, 2, 5, and 8 hours during induction, five females per time point were sampled and immediately snap-frozen in liquid nitrogen. Samples were stored at −80°C until processing.

### qPCR light/dark experiment

For the light/dark induction RT-qPCR experiment, 60 female anemones were divided into two groups. Whereas 30 females were induced for spawning under the same conditions as in the RNA-seq experiment, 30 females were induced in the same incubator that was covered to create dark conditions. In addition to a pre-induction sampling that served as a control for both groups, individual sea anemones from both treatment groups were sampled at the same time points as in the RNA-seq experiment at 1, 2, 5, and 8 hours during induction and frozen immediately. To ensure a proper spawning process, 87 females were induced by temperature and light and 48 by temperature in the dark; 81% and 77% of them, respectively, released egg sacks.

### RNA extraction

Total RNA was extracted from frozen anemone females using Quick-RNA MiniPrep Kit (Zymo Research, Irvine, USA), according to the manufacturer’s instructions with minor modifications. Lysis was performed using 3 mm glass beads in TissueLyser II (QIAGEN) and additional centrifugation at 1000 g for 1 min was then performed to precipitate unbroken particles. Genomic DNA residues were removed by Turbo DNase treatment (ThermoFisher Scientific) and final RNA was eluted in 80 μl nuclease-free ultra-pure water. Purified RNA concentrations were determined using NanoDrop 2000c spectrophotometer (Thermo Scientific). Extracted RNA from the RNA-seq experiment was quality assessed using TapeStation 2200 (Agilent Technologies) and extracted RNA from the qRT-PCR experiment was quality assessed using gel electrophoresis (1% agarose). RNA samples were stored at _80°C until use.

### Next-generation sequencing and data processing

Total RNA isolated from five females at each sampling point (RNA integrity number RIN > 8) was used for transcriptomic analysis. The 25 RNA samples were prepared using the Illumina TruSeq RNA Library Preparation Kit v3, according to the manufacturer’s protocol. The libraries were multiplexed on two lanes of an Illumina HiSeq2000 machine at the Life Sciences and Engineering Infrastructure Center at the Technion, Haifa. Illumina reads data (50 bp, single-end) were quality-filtered and adapter-trimmed using Trimmomatic 0.30 (90-96% mapping) (Bolger et al., 2014). Filtered reads were mapped and quantitated with STAR (version 2.4.2a) (Dobin et al., 2013) using the *Nematostella vectensis* genome (NCBI genome GCA_000209225.1). 7-13 million reads per sample were mapped uniquely to the genome assembly. Illumina results were deposited in the SRA database under accession no. PRJNA612671.

### Differential expression analysis

Differential expression analysis was conducted using generalized linear models (GLMs) in Bioconductor DESeq2 (in R 3.5.1), considering a single factor: induction time (Love et al., 2014). Differential expression was considered to be significant at p-value < 0.05 (adjusted using Benjamini-Hochberg correction), and transcript abundance is presented as FPKM (fragments per kilobase million) values produced in Deseq2 (Table S1). MA plots were generated with DEseq2 package in R language using DEseq2 lfcShrink function. Heatmaps of normalized FPKM values of all the significantly expressed transcripts were generated using the heatmap.2 function from package GPLOTS (v2.17.0) in R. Expected factor-dependent trends were verified using non-metric multidimensional scaling (nMDS) graphs in Vegan (https://github.com/vegandevs/vegan), using samples Bray-Curtis dissimilarity matrix.

### Gene Ontology (GO) enrichment analysis

Functional enrichment analysis was conducted using Bioconductor GoSeq (in R 3.5.1) (Young et al., 2010), with the Wallenius bias method, to allow correcting for gene-length biases in differential transcript expression. In addition, functional enrichment was detected using the score-based algorithm GSEA (Sergushichev, 2016; Subramanian et al., 2005) considering log_2_ fold changes as scores and GO terms were grouped to representative GOs using REVIGO (Supek et al., 2011). Non-canonical database for GoSeq was first built using Trinotate (Bryant et al., 2017). Heatmaps of significantly expressed transcripts within selected terms were generated using the heatmap.2 function from package GPLOTS (v2.17.0). In addition, networks and connections between transcripts were produced using KEGG pathway map (https://www.kegg.jp) (Kanehisa et al., 2016) and STRING (https://string-db.org/).

### Real-time qPCR

cDNA was synthesized using High Capacity cDNA Reverse Transcription Kit (Applied Biosystems) following manufacturer’s protocol without RNase inhibitor and modification of Step two in the PCR reaction program, which lasted 60 min. Primers were designed based on the exon-exon junction using Primer Express 3.0 software (Applied Biosystems) for a set of 10 transcripts of interest, which were significantly up-or down-regulated in the RNA-seq experiment (sequences were obtained from the published *Nematostella* JGI database https://mycocosm.jgi.doe.gov/Nemve1). Primer sequences used in this study are listed in Table S3.

The qPCR reaction was performed using StepOnePluse (Applied Biosystems) 96-well machine with Fast SYBR™ Green Master Mix (Applied Biosystems) as described before (Elran et al., 2014). The thermal profile was set to 95°C for 20 sec, followed by 40 amplification cycles of 95°C for 3 sec, 60°C for 30 sec, dissociation cycle of 95°C for 15 sec and 60°C for 1 min and then brought back to 95°C. *FUE* (far upstream element-binding protein 3) transcript (244532) was used as an internal reference control. The quantification analysis was done by the comparative CT method (Schmittgen and Livak, 2008), and all samples were quantified relatively to time 0 (pre-induction). Results are presented as the means with standard error of four biological replicates. While most data were normally distributed according to Shapiro-Wilk test, the rest were log_10_ transformed and retested for normal distribution. The significance of the results was determined using two-sample Student’s *t*-test or the nonparametric Mann-Whitney *U*-test for data that were not normally distrusted after transformation (SPSS software v. 20). The results were considered statistically significant if the null hypothesis could be rejected at the *P*<0.05 level.

## Results

### Transcriptomic analysis during spawning induction

To understand the molecular pathways leading to the release of oocytes in *Nematostella*, females were induced by temperature elevation and light exposure (see Methods) and then sampled for RNA-Seq analysis at 1, 2, 5, and 8 hours during induction in addition to a pre-induction control sampling (Fig. 1A). Transcriptome sequencing produced an average of 16.5 million raw reads per sample, filtered and mapped to the *Nematostella vectensis* genome (NCBI genome assembly GCA_000209225.1 ASM20922v1). Based on FPKM normalized read counts per transcript per sample, we estimated the distance between samples using non-Metric Multidimensional Scaling (nMDS) ordination, which indicated that the distance between the baseline and the treated samples grows with time (Fig. 1B). Generalized linear model in DESeq revealed ∼1,500 transcripts that were found to be differentially expressed (FDR <0.05) during spawning induction, as compared to pre-induction levels (Fig. 1C, Table S1). MA plots (Fig. 1D-G) demonstrate that during the induction process, both the number of differentially expressed transcripts and their fold-change magnitudes are gradually increased. This gradual effect is also seen in a heatmap, which additionally shows two distinct groups with opposite trends (Fig. 1C). The first group includes 608 transcripts that were upregulated eight hours after induction began, whereas the second group includes 628 transcripts that were downregulated eight hours into induction.

**Fig. 1:**
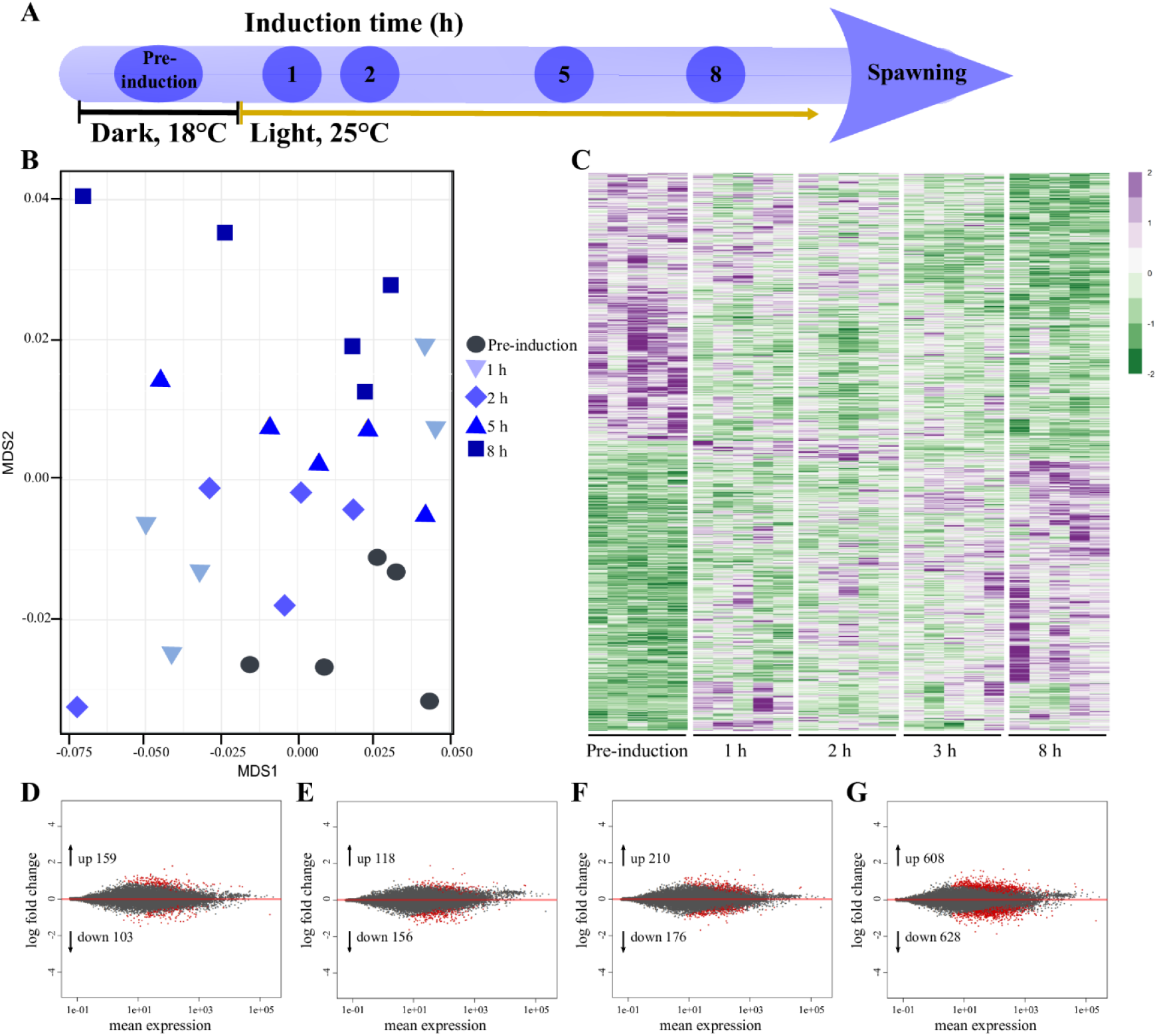
Transcriptome analysis of *Nematostella* females before and at different time points during spawning induction. (A) Schematic presentation of the transcriptome experiment. Induction time refers to time of temperature and light stimulus to successful spawning. Sampling time points are indicated in circles. Dark 18°C (pre-induction) and light 25°C (induction) regimes are indicated by black and yellow lines. (B) Non-metric, multidimensional scaling ordination indicates a time-dependent clustering of samples. Color-coded numbers on the right indicate sampling time points (n = 5 individuals per time point). (C) Heatmap of differential expressed genes at pre-induction and at the four sampling time points during induction. Rows represent transcripts, columns represent the tested individuals (n = 5). Expression level is indicated by z-score. The full list can be found in Table S1. (D-G) MA plots showing differentially expressed transcripts at 1 h (D), 2 h (E), 5 h (F) and 8 h (G) after induction began, compared to pre-induction. The numbers of up- and down-regulated transcripts are indicated within each plot. Y-axis represents the log_2_ fold change, and the X-axis represents mean normalized read counts. Red dots represent differentially expressed transcripts.

### Gene Ontology (GO) enrichment analysis

Next, we analyzed the enriched pathways of the differentially expressed transcripts during the experiment timeline. GO analysis revealed that 88% of the significantly enriched upregulated biological processes and molecular function categories were either specific to the first hour of induction (n=117, 42%) or shared between two or more time points (n=126, 46%) (Fig. 2, Table S2). Processes overrepresented among upregulated transcripts included terms related to light reception and chemical stimuli involving numerous receptors, as well as processes linked to the ECM and the nervous system. Among the significantly enriched downregulated biological processes and molecular function categories, 47% (n=168) were shared between time points and included terms related to cell cycle, oxidation-reduction and various metabolic processes (Fig. 2, Table S2).

**Fig. 2:**
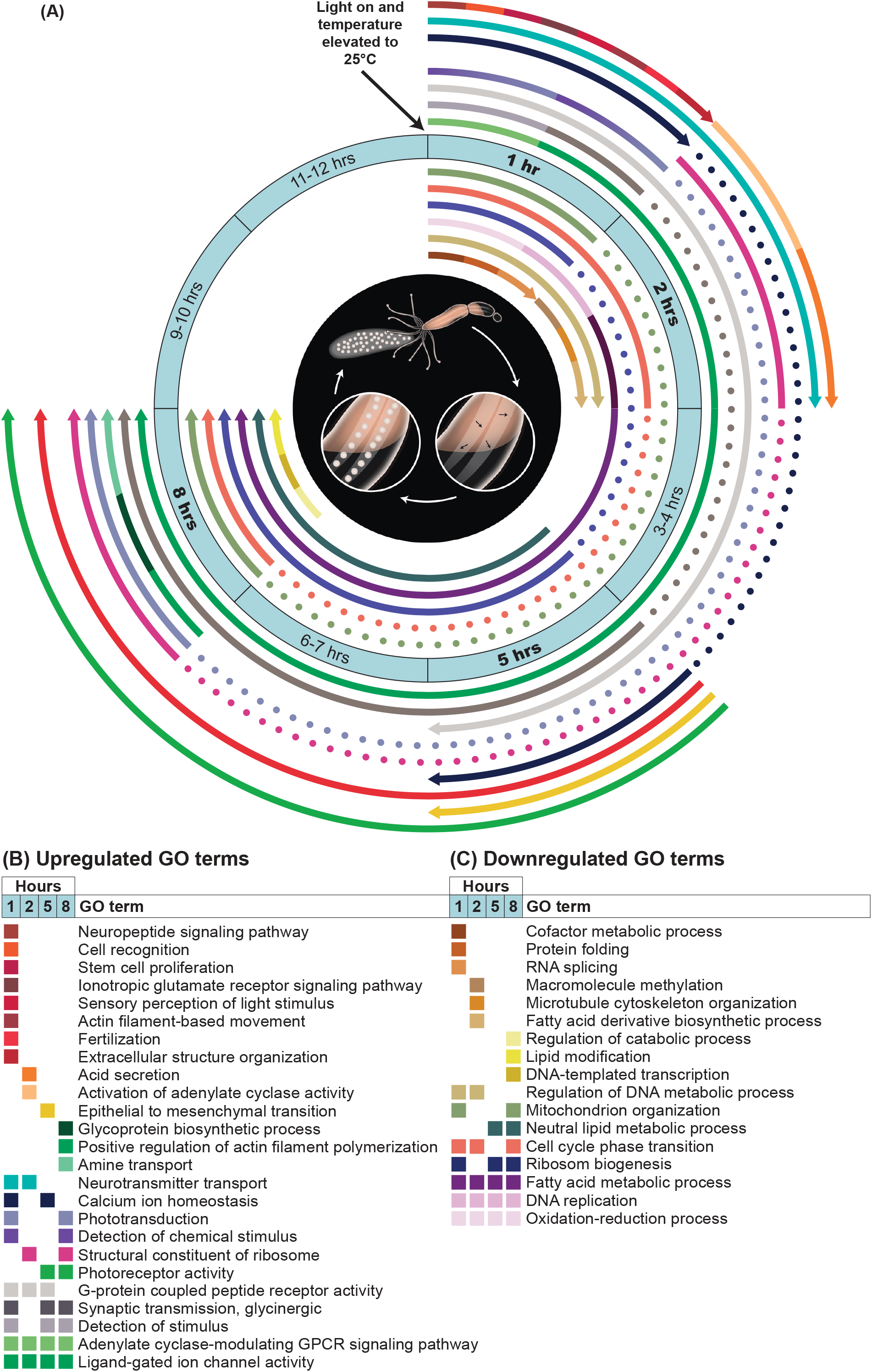
Infographic cartoon presenting temporal distribution from representative GO terms during spawning induction. (A) The middle circle indicates the timeline of induction, from light and temperature change to gamete release. Sampling time points are marked in bold. Each arc indicates the start point and endpoint (arrowhead) of up- or downregulated GO terms, which are shown in different colors; dashed arcs indicate temporary pauses in enrichment for the GO terms. Outer arcs represent upregulated enriched GO terms, and inner arcs represent enriched downregulated GO terms. Illustration in the inner black circle show the main stages of the induction process: oocytes develop in mesenteries, leading to the release of the egg sack into the water after approximately 12-15 hours. (B) Upregulated and (C) downregulated representative GO terms at 1 h, 2 h, 5 h and 8 h during spawning induction. The same color scheme as in A is used, and the color of each term is annotated. The complete list of all GO terms can be found in Table S2

### Processes related to light, ECM, and cytoskeleton are upregulated during spawning induction

Since light is one of the triggers of spawning induction in *Nematostella* (Fritzenwanker and Technau, 2002), we analyzed the GO term sensory perception of light stimulus (GO:0050953), which was upregulated during the first hour of induction (Fig. 2, 3A). The analysis revealed that transcripts associated with this term are expressed mainly at one hour and eight hours into induction and that many are receptors, specifically G protein-coupled receptors (GPCRs) (Fig. 3A). Moreover, five and eight hours into induction, there was an upregulation of transcripts involved in photoreceptor activity (GO:0009881) (Fig. 2, 3B), and eight hours into induction, there was also an upregulation of the photoperiodism process (GO:0009648), though not upholding significance (*p*=0.05). Together, these two processes regulated 11 transcripts, five of which are core components of the circadian clock. Among the circadian clock transcripts, *Clock* and three cryptochromes (*Cry1a, Cry1b* and *Cry*) were downregulated before induction and their expression elevated during the process, peaking after eight hours. The transcript *Cry1* displayed an opposite expression trend, as it was upregulated before induction started and decreases during the process (Fig. 3B). Other transcripts that are related to photoreceptor activity, such as melanopsin, rhodopsin, protease usp6 n-terminal-like, rhodopsin-like GPCR and opsin BCRH2-like, displayed a similar expression trend of upregulation during the induction process.

**Fig. 3:**
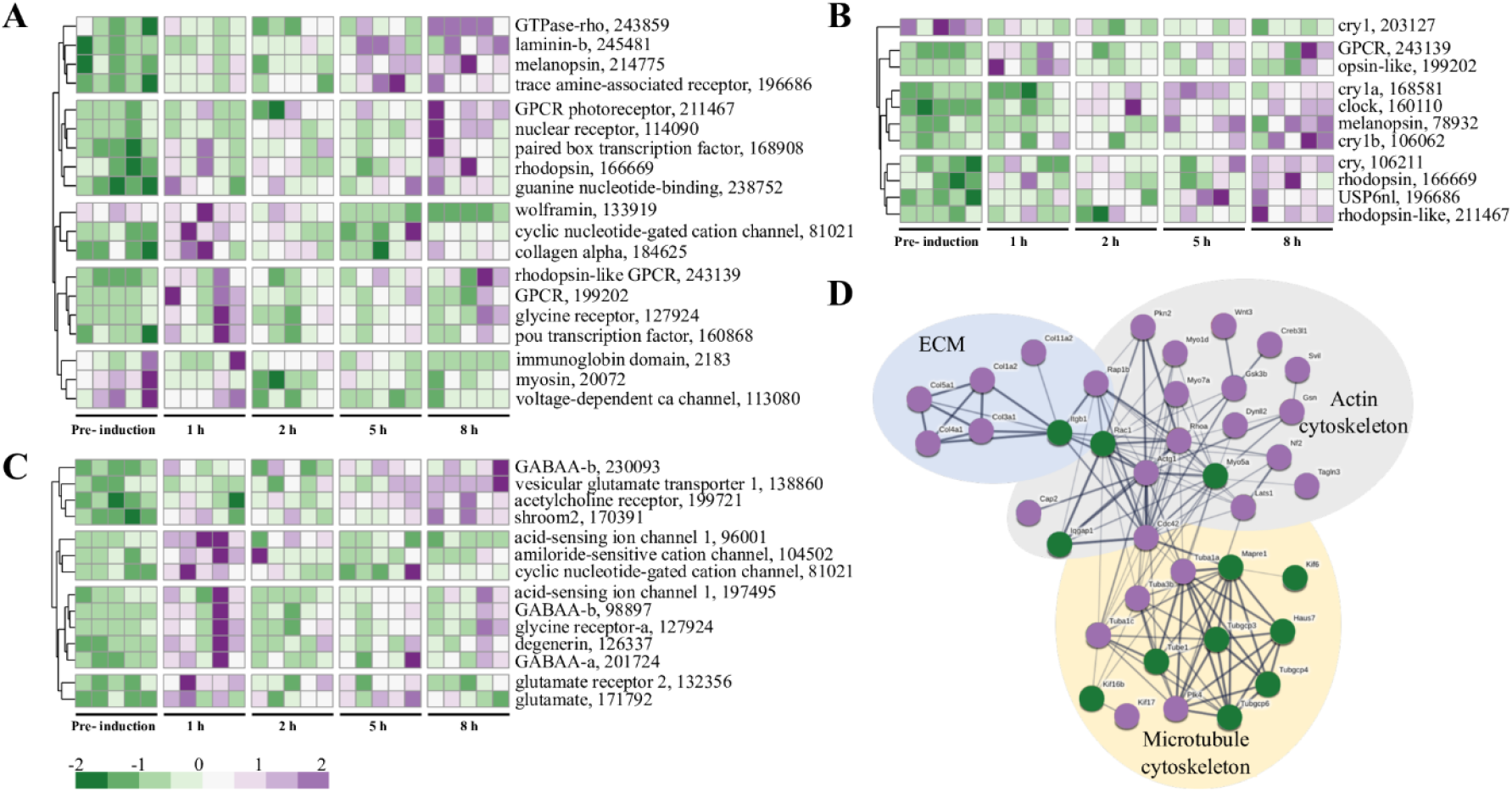
Analysis of transcripts related to light, ion channel, ECM and cytoskeleton. Heatmap of transcript expression related to (A) sensory perception of light stimulus (GO:0050953), (B) photoreceptor activity (GO:0009881) and photoperiodism (GO:0009648) and (C) ligand-gated ion channel activity (GO:0015276) at the different sampling time points during induction, relative to the pre-induction control (FDR<0.05). Rows represent transcripts, columns represent the tested individuals (n = 5 per time point). Expression level is indicated by z-score. Numbers are JGI IDs. (D) STRING visualization of protein interactions of cytoskeleton- and ECM-related transcripts. Upregulated transcripts are in purple and downregulated in green. The proteins presented in the network are mouse homologs. Thickness of the connecting lines indicates the strength of data support.

We also analyzed the term ligand-gated ion channel activity (GO:0015276), which was common to all sampling time points (Fig. 2, 3C). The analysis revealed four different GABA_A_ receptor subunits and three transcripts related to the excitatory neurotransmitter glutamate. Two of the glutamate-related transcripts (171792, 132356) displayed similar patterns of upregulation, mainly at the first hour of induction, whereas the third transcript (138860) was strongly upregulated at the eighth hour of induction (Fig. 3C).

We then used STRING to analyze the cytoskeleton and ECM proteins related to the GO terms actin filament-based movement (GO:0030048), extracellular matrix structural constituent (GO:0005201), and positive regulation of actin filament polymerization (GO:0030838), which were upregulated at the first, second, and eighth hour of the induction process, respectively. The analysis revealed a complex network that can be divided into three main groups: ECM, actin cytoskeleton and microtubule cytoskeleton (Fig. 3D). Some of the microtubule isoforms were upregulated, whereas others were downregulated and displayed a weak connection to isoforms of motor proteins, such as dynein and kinesin (Fig. 3D). The myosin motor proteins are connected to actin, which showed strong connections to three GTPases: *RhoA, Cdc42*, and *Rac1*. While the first two were upregulated, *Rac1* was downregulated. Additionally, connected to actin is the integrin receptor *Itgb1*, which was downregulated and strongly connected to different isoforms of collagens comprising the ECM network, all found to be upregulated (Fig. 3D). Overall, the upregulated processes indicate that spawning induction is perceived by different receptors, some of which are light-sensitive receptors that initiate signaling cascades and cytoskeletal rearrangement.

### Metabolic processes are downregulated during spawning induction

A large group of downregulated processes included seven terms associated with metabolism. Only three of the processes were shared among time points, two of which are related to lipid metabolism, namely fatty acid metabolic process and neutral lipid metabolic process (Fig. 2). KEGG analysis of the term fatty acid metabolic process (GO:0006631) revealed that nine transcripts function in fatty acid degradation, and except one, all of them were downregulated (Fig. S1). These results indicate that degradation of fatty acid and consequently the formation of acetyl-CoA, which is required for ATP production, are reduced during spawning induction.

### Light effect on the induction process

Next, we performed real-time qPCR on selected transcripts obtained from RNA-seq to test their transcriptional response to light during induction. For that, *Nematostella* females were induced by temperature elevation under either light or dark conditions and were sampled at the same time points as in the RNA-seq experiment (Fig. S2A). While in both groups spawning rates were about 80%, some transcripts displayed different expression patterns in response to light (Fig. S2). Out of seven clock-related transcripts examined, five transcripts (*Cry1a, Cry1b, Clock*, BHLH transcription factor *Helt*, and glutamate receptor *Grin1*) were upregulated during light induction and at the eighth hour of the dark induction, whereas one transcript, thyrotroph embryonic factor (*Tef*), was downregulated as induction proceeded under both light and dark conditions (Fig. S2B-H). We additionally tested two cytoskeleton-related transcripts, tubulin 1 (*Tuba1*) and *Wnt3*, and the aquaporin (*Aqp*) water-selective channel. With the exception of *Aqp*, no significant differences in expression levels were found between the two treatment groups (Fig. S2I-K). These results support the findings of light perception during spawning induction but also demonstrate that temperature raise is a stronger signal for spawning.

## Discussion

To explore the molecular mechanisms and pathways underlying spawning induction in the basal metazoan *Nematostella*, we analyzed the transcriptome of females before and during the induction process. Our results revealed a diverse array of up-and-downregulated processes, which lead to successful spawning. Furthermore, a complementary light/dark induction experiment revealed that the expression levels of most tested transcripts changed significantly under both conditions, demonstrating that temperature change is a stronger signal for spawning. These molecular data support the predominance of temperature change over light, which was shown before in spawning experiments (Fritzenwanker and Technau, 2002).

In cnidarians, light has been shown to have a fundamental role in regulating sexual reproduction. The spawning of the hydromedusa *Clytia hemisphaerica* and *Cladonema pacificum* is triggered by light-dark transition and is mediated by opsin photoreceptors from the GPCR family (Lamb et al., 2007; Quiroga Artigas et al., 2018; Quiroga Artigas et al., 2020). In the corals *Acropora millepora* and *Acropora digitifera*, the photopigment melanopsin, which belongs to the opsin family, is upregulated during the spawning night, possibly triggering different signaling cascades contributing to gamete release (Kaniewska et al., 2015; Rosenberg et al., 2017). We found upregulation of several GO terms related to light reception and transduction as well as to GPCR and ion channel activity, including melanopsin and rhodopsin receptors from the GPCR family and a glutamate receptor. This suggests that in *Nematostella*, light-sensitive proteins are responsible for receiving the light signal and initiate different cascades, such as GPCR signaling and glutamate release. Additionally, our qPCR analysis showed that the glutamate receptor *Grin1*, an ion-gated channel (Lau and Zukin, 2007; Paoletti et al., 2013), was upregulated at the eighth hour of both light and dark inductions. Previous studies have shown that the glutamate pathway interacts closely with the molecular feedback loops of the circadian clock (Ikeda, 2004; Ikeda et al., 2003). In *A. digitifera* and *A. millepora*, upregulation of transcripts related to glutamate release by photoreceptors was observed during the spawning event (Kaniewska et al., 2015; Rosenberg et al., 2017). Together, these results suggest a conserved role for GPCR and glutamate signal transduction during the reproduction process of *Nematostella*. However, the exact mechanism is yet to be uncovered.

The core components of the circadian clock, cryptochromes and *Clock*, have been identified in *Nematostella* and were shown to have diel rhythmicity. Moreover, there is evidence that *Nematostella* exhibits circadian behavior and physiology under diel light cycles that are maintained upon the removal of light (Hendricks et al., 2012; Oren et al., 2015; Reitzel et al., 2010; Reitzel et al., 2013). In *A. millepora*, two cryptochromes that displayed rhythmic gene expression under light:dark cycles were suggested to play a role in synchronizing spawning (Levy et al., 2007). Our results showed that the expression of *Cry1a, Cry1b* and *Clock* was upregulated during light induction, while in the dark induction they were upregulated at the eighth hour only. Since cultivated cultures of *Nematostella* can be induced in light and dark conditions, our results suggest that *Nematostella Cry*s’ and *Clock* are not essential for initiating the process, yet, may play non-circadian roles later during spawning induction. Another clock-related gene, *Tef*, which belongs to the PAR family of transcription factors, was shown to be expressed in vertebrates with a circadian rhythms (Fonjallaz et al., 1996) and to regulate downstream light-induced genes under circadian control in zebrafish (Gavriouchkina et al., 2010). Our results demonstrated that in *Nematostella, Tef* is not affected by light, as it was downregulated in both light and dark conditions. This is in line with the findings in *A. millepora*, where *Tef* was downregulated prior to and during spawning (Kaniewska et al., 2015). It is possible that *Tef* functions as a repressor of a yet unknown pathway and therefore its downregulation allows the pathway to operate. Although the circadian clock plays an important role in the reproduction of many organisms from different phyla (Beaver et al., 2002; Boden et al., 2013; Giebultowicz et al., 1989; Naylor, 2010), our results indicate that spawning of *Nematostella*, during controlled laboratory inductions, is most likely not regulated by the endogenous clock. Notably, the circadian expression pattern of these clock genes is strongly dependent on consistent light:dark cycles (Leach and Reitzel, 2019; Reitzel et al., 2010); however, the studied sea anemones were maintained in constant temperature and darkness for a long period and were not entrained by daily exogenous cues. Therefore, more investigation is required to understand the role of circadian clock components during spawning induction in *Nematostella*

One of the expected changes during spawning induction is the rearrangement of cytoskeletal networks. In the corals *Acropora gemmifera, A. millepora* and *A. digifera*, cytoskeletal reorganization was elevated during and after spawning (Kaniewska et al., 2015; Oldach et al., 2017; Rosenberg et al., 2017). Our results demonstrated that likewise, extensive changes in the cytoskeleton and its related proteins occur during spawning induction in *Nematostella*. We have discovered transcripts that are involved in actin regulation, such as the small GTPases *RhoA, Rac1*, and *Cdc42* (Fig. 4). *Cdc42* promotes the formation of filopodia (Nobes and Hall, 1995) and *RhoA* regulates the formation of linear actin network, focal adhesion and stress fibers, which are the major contractile structures required for cell adhesion during migration (Etienne-Manneville, 2004; Parri and Chiarugi, 2010). When *RhoA* is activated, it affects the actomyosin contractile network (Piekny et al., 2005). This stands in line with our results showing upregulation of *RhoA* and myosins. *Rac1* controls the formation of branched actin network and the formation of lamellipodia in the leading edge of motile cells (Parri and Chiarugi, 2010). *RhoA* and *Rac1* usually antagonize each other (Guilluy et al., 2011) and the balance between them was shown to control cell shape (Chauhan et al., 2011). The finding of upregulation of *RhoA* and downregulation of *Rac1* suggests that such balance may allow the mesenteries to change shape and create space for oocyte development. Additionally, it was shown that in migrating cells, branched actin network is localized in the leading edge of the cell, while stress fibers are present in the cell body, anchoring it to the surface through focal adhesions (Friedl and Wolf, 2003). Our findings indicate a similar actin organization, suggesting that cell migration occurs during spawning induction.

**Fig. 4:**
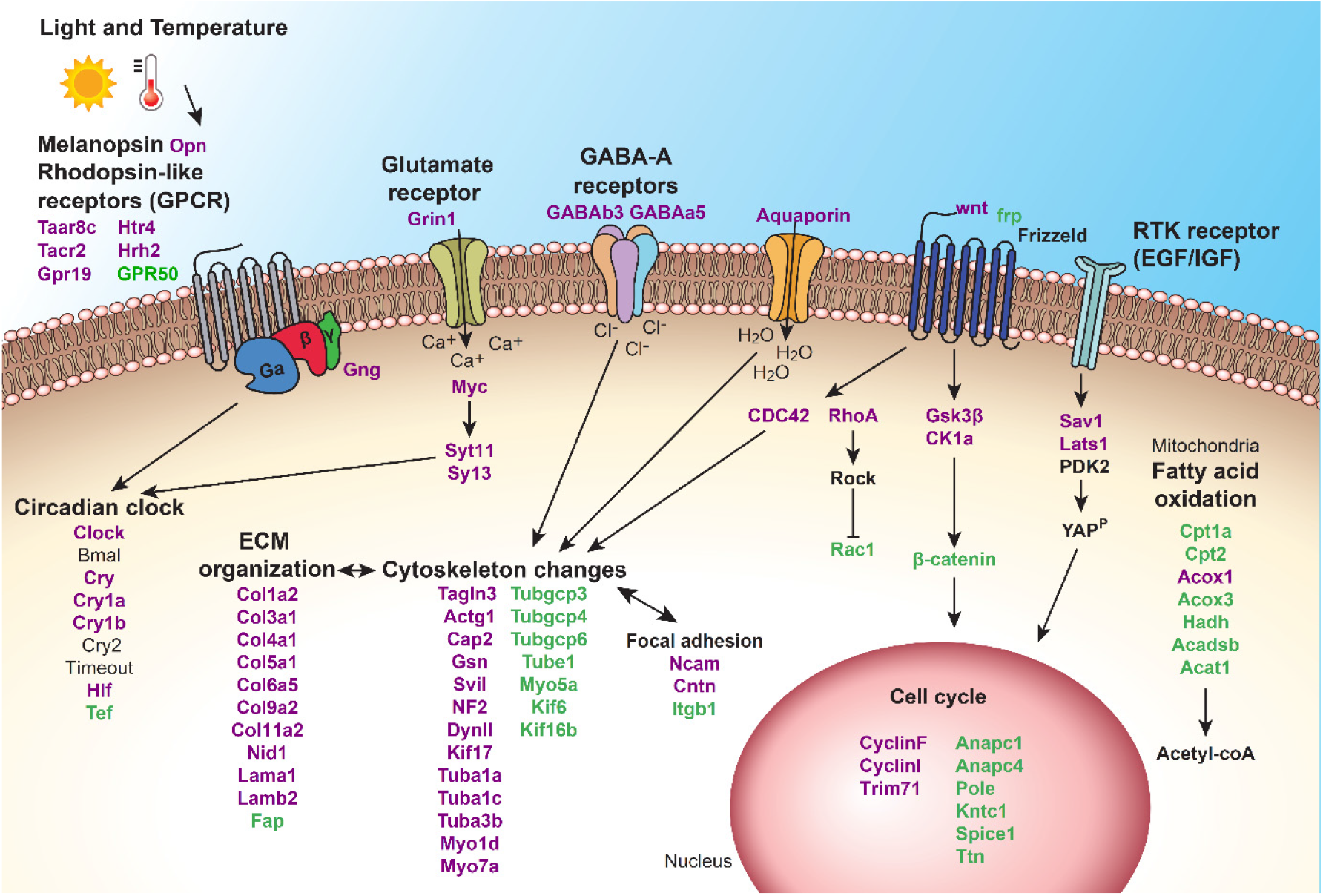
Proposed model for molecular signaling during spawning induction in *Nematostella vectensis*. The process is triggered by light and temperature elevation. We propose that light stimulates melanopsin-like homologue, which activates rhodopsin and glutamate receptors and initiates GPCR signaling cascades. Other receptors are also activated and together they affect cell cycle, cytoskeleton and ECM organization, focal adhesion, fatty acid oxidation and circadian clock components. Eventually, these processes lead to successful spawning. Upregulated transcripts are written in purple and downregulated transcripts are in green.

The ECM serves as a substrate for cell migration and transfers signals into the cell from the surrounding environment through adhesion receptors such as integrins, which promote reorganization of the cytoskeleton (Damsky and Werb, 1992; Hynes, 2002). In addition, ECM remodeling plays a crucial part in regulating diverse cellular processes such as cell shape and growth (Bonnans et al., 2014). Our results showed upregulation of many different isoforms of collagen, the most abundant ECM proteins (Karsenty and Park, 1995) (Fig. 4), possibly indicating ECM rearrangement.

Many of the downregulated GO terms we found are related to metabolic processes, such as fatty acid metabolism. This could imply that during induction, less energy is generated in the mesenteries. In contrast, fatty acid oxidation is increased in the oocytes (Lotan et al., 2014). We additionally found that transcripts related to cell cycle, an energy consuming process, were downregulated (Fig. 4). Furthermore, DNA, RNA and protein related processes were downregulated, suggesting that cell maintenance was reduced. Altogether, these results indicate that there is a decrease in metabolic processes that are not required for spawning. The mesenteries were previously shown to host the synthesis of the fatty acid-rich vitellogenin (Vtg), which is the main yolk protein in *Nematostella* oocytes (Levitan et al., 2015; Lotan et al., 2014). In fish, vitellogenesis is accompanied by an increase in fatty acid in the liver, where the process occurs. Decrease in fatty acid in the liver and their increase in gonads were observed before spawning and occurred due to transport of Vtg to the oocyte (Blanchard et al., 2005; Huynh et al., 2007). We, therefore, postulate that the observed decrease in fatty acid breakdown can be a result of Vtg synthesis. Together, these findings indicate that the mesenteries function as gonads during spawning in *Nematostella*.

To conclude, this study reveals molecular pathways underlying spawning induction in female sea anemones, *Nematostella vectensis*. Our results demonstrate that upon the reception of light and temperature signals, processes involving numerous receptors, circadian clock components, the cytoskeleton and ECM are upregulated, whereas housekeeping processes are mostly downregulated. As shown by the conceptual model in Figure 4, we suggest that these changes may enable the role of the mesenteries as a gonad-like tissue for the developing oocyte and eventually lead to spawning. Our findings demystify the progression of spawning induction in basal metazoan and could assist in deeper understanding the evolution of spawning patterns of higher organisms that share a common ancestor with Cnidaria.

## Acknowledgments

We thank the Bioinformatics Service Unit at the University of Haifa and specifically M. Lalzar for her assistance. This study represents partial fulfillment of the requirements for a Ph.D. thesis for M. Rinsky at the Faculty of Life Sciences Bar-Ilan University, Israel.

## Author Contributions

S.R. designed and performed the experiments with the assistance of V.B. M.R., S.R, O.L., and T.L. analyzed the data, A.M. performed the bioinformatics, O.L. and T.L. conceived and supervised the project, M.R. wrote the manuscript with input from all authors. All authors discussed the results and commented on the manuscript. All authors read and approved the final version of the manuscript.

## Data Availability

The sequencing data generated during this study has been deposited to the Sequence Read Archive (SRA), under accession PRJNA612671.

## Supplementary Figures and Tables

**Table S1:** Transcriptome analysis of all transcripts during spawning induction (Excel file).

**Table S2:** Enrichment of up- and down-regulated GO terms at 1, 2, 5 and 8 h during induction (Excel file).

**Table S3:** List of primers used in qPCR expression analysis.

**Fig. S1:** KEGG metabolic map of fatty acid degradation.

**Fig. S2:** Real-time qPCR analysis before and during light or dark induction.

